# Antiviral sugar baits to reduce arbovirus transmission: a proof-of-concept study

**DOI:** 10.64898/2026.09.07.749591

**Authors:** Maeva Van Langendonck, Ana L. Rosales-Rosas, Shannan-Leigh Macleod, Johan Neyts, Fernando Genta, Leen Delang

## Abstract

Dengue virus (DENV) and chikungunya virus (CHIKV) remain major global health threats, yet approved antiviral therapies are lacking. Antiviral sugar baits (AVSBs) represent a novel transmission-blocking strategy that exploits the natural sugar-feeding behaviour of mosquitoes to deliver antiviral compounds directly to the mosquito vector. In this study, we evaluated the antiviral efficacy of AVSBs containing JNJ-A07, β-D-N4-hydroxycytidine (NHC, EIDD-1931), molnupiravir (MPV), and 4′-fluorouridine (4′FlU) in *Aedes aegypti* mosquitoes. An *ex vivo* mosquito gut model was used to select antiviral concentrations and to assess the ability of 4′FlU to reach the mosquito midgut following sugar feeding. AVSBs were subsequently evaluated for their effects on mosquito attractiveness, longevity, fecundity, and fertility, as well as their ability to suppress DENV and CHIKV infection *in vivo*. 4′FlU potently inhibited CHIKV replication in *ex vivo* mosquito guts and retained antiviral activity following sugar-bait administration. None of the antiviral compounds affected mosquito attraction to the bait, while only modest and compound-specific effects on mosquito fitness were observed. AVSBs containing JNJ-A07 significantly reduced DENV infection and dissemination, whereas 4′FlU-containing AVSBs significantly reduced CHIKV infection and viral loads. In contrast, NHC- and MPV-containing AVSBs did not exhibit antiviral activity *in vivo*. Collectively, these findings provide proof-of-concept for AVSBs as a novel strategy to reduce arbovirus transmission by targeting viral replication within the mosquito vector, suggesting that AVSBs could complement existing arbovirus control measures.

## Introduction

Mosquito-borne viruses, in particular dengue virus (DENV) and chikungunya virus (CHIKV), pose a significant threat to global public health due to their widespread transmission, substantial disease burden in humans, and potential to cause severe clinical consequences in some infected patients [1]. DENV is classified within the genus *Orthoflavivirus* of the *Flaviviridae* family. It is the most widespread mosquito-borne virus, with an estimated 390 million infections annually. The virus is currently endemic in more than 100 countries, mainly in tropical and subtropical regions of the Americas, Asia, Africa, and Australia [2]. CHIKV belongs to the *Alphavirus* genus of the *Togaviridae* family. The incidence of CHIKV has increased dramatically over the past few decades, with major outbreaks reported in more than 100 countries across Africa, Asia, the Indian subcontinent, and the Americas [3]. Both viruses are mainly transmitted by the bite of *Aedes (Ae.) aegypti* and *Ae. albopictus* mosquitoes. Infection with DENV or CHIKV manifests as an acute febrile illness characterized mainly by polyarthralgia and myalgia. A notable proportion of DENV-infected patients (ranging from 0.5% to 5%) can develop severe dengue, reaching a fatality rate exceeding 20% when supportive care is absent. On the other hand, 25 to 75% of CHIKV patients will develop debilitating chronic arthralgia, which can persist for several months or years following the initial infection [4]-[5].

Despite the growing global burden of mosquito-borne viruses, the use of licensed vaccines including Qdenga [6] for dengue and Vimkunya [7] for chikungunya, remains constrained. Furthermore, no approved antiviral therapies currently exist for either disease [6]. Over recent years, several antiviral molecules effective against arboviruses have been identified. Relevant to this study are JNJ-A07, molnupiravir (MPV), β-d-N4-hydroxycytidine (NHC), and 4’Fluorouridine (4′FlU). JNJ-A07, a highly potent DENV NS4B inhibitor, has demonstrated antiviral efficacy *in vitro* and in mouse studies [8]. NHC is the active metabolite of the oral prodrug MPV. NHC has demonstrated potent anti-CHIKV activity *in vitro* and in mouse models as a lethal mutagen that impairs viral replication [9]-[10]. Likewise, the ribonucleoside analogue 4′FlU has demonstrated potent anti-CHIKV activity *in vitro* and in mouse models, reducing viral burden, tissue swelling, and inflammation [11].

Although considerable efforts have focused on developing antivirals for therapeutic use in humans, exploiting antiviral activity within the mosquito vector represents a novel approach to interrupt virus transmission. Mosquitoes could acquire antiviral drugs through a blood meal from a drug-treated individual or through tarsal exposure (via the legs) when resting on drug-treated surfaces, similar to the uptake of insecticides from long-lasting insecticidal nets [12]. Such transmission-blocking strategies exploit mosquito behaviors, such as blood feeding or resting behavior, and existing vector-control measures, to deliver small molecules that interrupt pathogen replication within the vector and subsequently transmission to humans.

Previous studies have demonstrated the feasibility of this concept. For instance, tarsal exposure to the antimalarial drug atovaquone effectively blocked *Plasmodium* development in *Anopheles* mosquitoes [13] and was later shown to reduce CHIKV transmission by *Ae. aegypti* mosquitoes [14]. In addition, JNJ-A07 was shown to reduce DENV infection, dissemination, and transmission in *Ae. aegypti* mosquitoes when delivered through a blood meal [15]. In contrast, while NHC effectively inhibited CHIKV replication in *ex vivo* cultured *Ae. aegypti* mosquito guts, the antiviral activity was not maintained following delivery through a blood meal [16]. Collectively, these studies demonstrated that antiviral compounds can have antiviral activity within mosquito tissues, but their efficacy may depend strongly on the delivery route and pharmacokinetic properties of the compound.

Beyond blood-feeding and tarsal exposure, antiviral drugs could be delivered through the mosquito’s natural sugar feeding behavior. Attractive targeted sugar baits (ATSBs) exploit the frequent sugar feeding of adult mosquitoes to deliver active compounds and have been traditionally used for insecticide-based vector control [17]. This approach has also been successfully adapted to deliver other molecules, such as short hairpin RNA targeting essential mosquito genes, causing high mortality rates [18]. Replacing the insecticidal component with an antiviral drug could therefore provide a complementary strategy to interfere with arbovirus replication within the mosquito. However, the potential of antiviral sugar baits has not yet been explored.

Building on our previous findings demonstrating that antiviral compounds can be delivered through a blood meal to inhibit arbovirus replication in mosquitoes, we investigated whether a similar antiviral effect could be achieved via sugar feeding. Because the uptake, processing, and metabolism of sugar meals differ substantially from those of blood meals, the effectiveness of antiviral delivery through this route remains unknown. Therefore, we evaluated whether antiviral sugar baits (AVSBs) could deliver antiviral compounds to *Ae. aegypti* mosquitoes and suppress DENV and CHIKV replication. By exploiting the mosquito’s natural sugar-feeding behavior, this study provides proof-of-concept for a novel transmission-blocking strategy.

## Materials and methods

### 1. Viruses

The CHIKV Indian Ocean strain 899 (GenBank accession no. FJ959103.1) was kindly provided by Prof. Drosten (University of Bonn, Bonn, Germany) [19]; DENV serotype 2 (DENV-2/TH/1974, isolated in 1974 from human serum collected in Bangkok, Thailand; GenBank MK268692.1) was kindly provided by Prof. Failloux (Institut Pasteur, France) [20]. Virus stocks were prepared by passaging the isolates on C6/36 cells, and viral titers were determined by performing plaque assays on Vero cells and BHK cells, respectively.

### 2. Compounds

JNJ-A07 was obtained from Janssen Pharmaceutica (Beerse, Belgium). The compound was dissolved in 100% DMSO at a 100 mM stock concentration, stored at 4°C. β-D-N4 - hydroxycytidine (NHC or EIDD-1931), molnupiravir (MPV or EIDD-2801) and 4’Fluorouridine (4’FlU) were purchased from MedChemExpress (Monmouth Junctions, NJ, USA) and dissolved in 100% DMSO at a 40 mM or 100 mM concentration, which was stored at −20 °C and protected from light until use.

### 3. *Aedes aegypti* mosquitoes

Eggs belonging to the *Ae. aegypti* Paea strain (Papeete, Tahiti, 1994) were obtained from Prof. Failloux via the Infravec2 consortium. For rearing, eggs were hatched in dechlorinated tap water, and larvae were transferred to plastic trays containing 3 L of water and fed daily with a yeast tablet (Gayelord Hauser, Saint-Genis-Laval, France) until pupation. Pupae were placed in small bowls and allowed to emerge inside 30×30×30 cm BugDorm cages (MegaView Science Co., Ltd., Taichung, Taiwan). Adult mosquitoes were provided with cotton balls soaked in a 10% sugar solution. The trays and cages were incubated at 28±1°C and 80% relative humidity (RH) with a light:dark cycle of 16:8 h.

### 4. Preparation of the AVSB

The sugar baits were prepared in a fresh 10% sucrose solution. Final DMSO concentrations in the AVSB were maintained below 0.5%. Sugar baits were administered via cotton balls soaked in the respective solutions.

### 5. *Ex vivo* mosquito gut assay with 4’Fluorouridine

#### Condition 1: Antiviral assay in ex vivo gut cultures

The mosquito *ex vivo* gut assay was performed as described previously [21]. In brief, dissected guts were infected with CHIKV 899 (1 × 10^4^ PFU/mL) for 2h at 28°C without CO_2_, washed three times with PBS and subsequently incubated in gut medium containing either DMSO (control) or 4’FlU (treatment). After 3 days of incubation at 28°C without CO_2_, individual guts were collected, homogenized, filtered, and assessed using end-point titration assays (Fig. 1).

**Fig 1.**
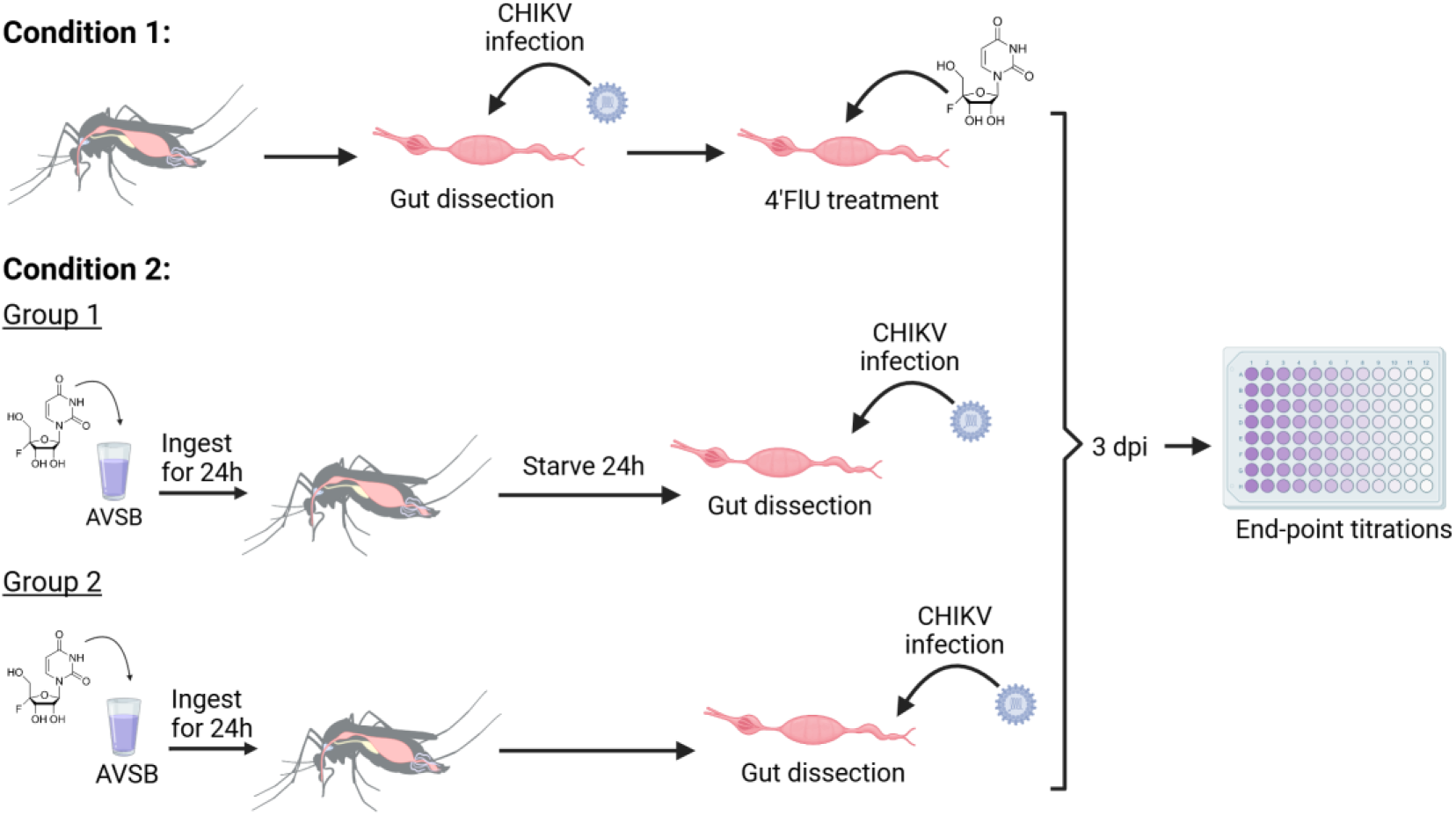
Schematic overview of the experimental set-up used to evaluate the antiviral efficacy of 4′-fluorouridine (4′-FlU) against CHIKV in *ex vivo Aedes aegypti* guts. Direct treatment of infected guts with 4′-FlU (Condition 1) was compared with delivery through antiviral sugar baits (AVSBs) (Condition 2) prior to gut dissection and *ex vivo* infection. In Condition 2, mosquitoes were either dissected immediately after 24h of AVSB feeding (Group 2) or after an additional 24h starvation period (Group 1). Viral titers in the guts were determined by endpoint titration at 3 days post-infection.

#### Condition 2: Prophylactic antiviral treatment

Female mosquitoes (5 – 7 days old) were placed in paper cups and starved for 24h prior to being offered either 200 µM 4’FlU sugar bait (treatment group) or a DMSO-containing sugar bait (control group) (prepared as described above), *ad libitum*. Following 24h of access to the sugar bait, mosquitoes in group 1 were starved for an additional 24h before gut dissection. In contrast, mosquitoes in group 2 were dissected immediately after the 24h sugar bait exposure period (Fig. 1). Dissected guts from both groups were subsequently infected *ex vivo,* maintained in fresh gut medium for 3 days. Hereafter, individual guts were collected, homogenized, filtered, and assessed using end-point titration assays.

### 6. Olfactory Attraction experiment

Female mosquitoes (5 – 10 days old, 40 per group) were aspirated and placed in a small cage (BugDorm). Mosquitoes were starved for 24h. Sugar baits containing compound or DMSO (control) were prepared in a 10% sucrose solution at a concentration of 20 µM for JNJ-A07, 100 µM for MPV and NHC, and 200 µM for 4’FlU. Cotton balls were soaked in the AVSB or control sugar bait and placed on opposite sides of the cage. Mosquito landings on each cotton ball were recorded for three constitutive observation periods of 10 minutes. A preference index (PI), using formula: 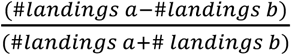, was calculated to assess whether the mosquitoes showed a preference for the antiviral or control sugar bait solution.

### 7. Longevity assay

Mosquitoes (5 days old) were placed in cups in groups of 20 (males and females separated) and sugar deprived for 24h. Sugar baits containing compound (treatment group) or DMSO (control group) were prepared in a 10% sucrose solution at a concentration of 100 µM for MPV and NHC, and 10 µM for 4’FlU. The mosquitoes were offered the sugar bait *ad libitum* for 7 days via soaked cotton balls. The mortality was recorded daily for 30 days.

### 8. Fecundity and fertility

Female mosquitoes (5 – 10 days old) were starved for 24h before receiving a freshly prepared antiviral (treatment) or DMSO (control) sugar bait solution, which was administered on a cotton ball for a period of 7 days. After 7 days, the mosquitoes were starved for 20h before receiving a blood meal, consisting of fresh rabbit erythrocytes supplemented with ATP (5mM), via the Hemotek feeding system. Mosquitoes were blood fed for 45 min. Thereafter, only fully engorged females were selected and incubated at 28°C with access to 10% sucrose for the remainder of the experiment. For the fecundity measurement, the ovaries of each mosquito were dissected on day 4 post blood feeding. The number of developed eggs per female were counted. For the fertility assay, the blood fed mosquitoes were individually placed in a cardboard cup containing an oviposition cup with a damp substrate (Whatman filter paper). The laid eggs were counted for each female, and the oviposition papers were individually hatched. The number of larvae that emerged from each egg paper was counted 3 days after hatching the eggs.

### 9. Mosquito exposure to antiviral sugar baits and virus oral infection

Female mosquitoes (3 – 5 days old) were starved for 20h (Fig. 2). Starved mosquitoes were offered the sugar bait solution containing one antiviral drug (JNJ-A07 at 2 µM or 20 µM; MPV at 100 µM; NHC at 100 µM or 4’FlU at 200 µM) or DMSO (control) for 2 days via a cotton ball. On day 3, the antiviral sugar bait was removed, and the mosquitoes were starved for an additional 24h.

**Fig. 2.**
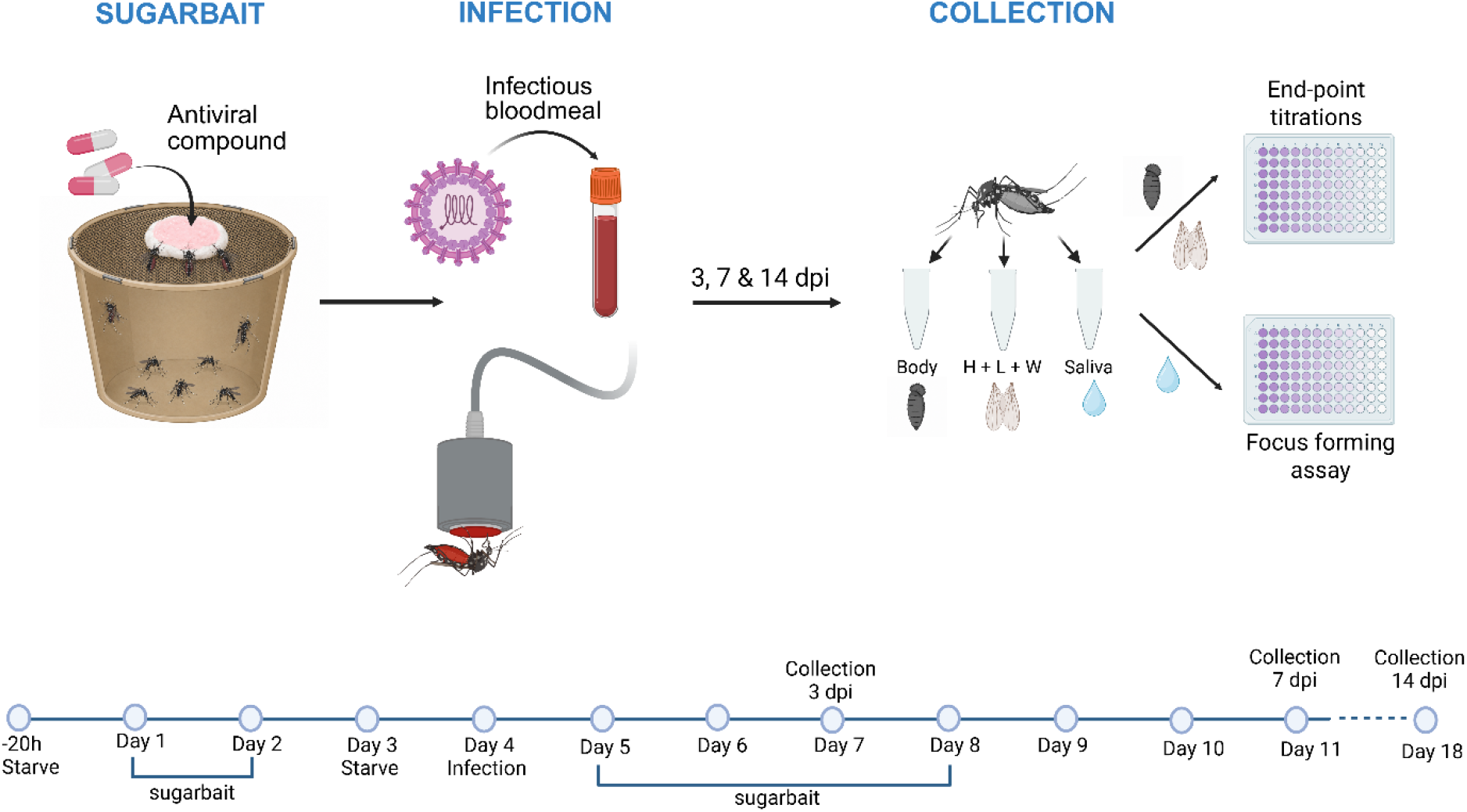
Experimental workflow for antiviral sugar bait treatment and virus infection in mosquitoes. Mosquitoes were exposed to antiviral sugar baits prior to and following DENV-2 or CHIKV infection. Saliva, bodies, and heads with wings and legs (H + L + W) were collected at 3, 7, or 14 dpi and analyzed for infectious virus.

Next, the mosquitoes were offered an infectious blood meal with either DENV-2 (JNJ-A07-treatment) or CHIKV (MPV, NHC, and 4’FlU-treatment). The blood meal consisted of fresh rabbit erythrocytes supplemented with ATP (5mM), FBS, and virus (DENV-2, 3 × 10^7^ PFU/mL or CHIKV, 1 × 10^7^ PFU/mL), using the Hemotek feeding system. Mosquitoes were fed for 45 minutes, whereafter fully engorged mosquitoes were sorted while cold anesthetized and maintained at 28°C for the remainder of the experiment. After the infection, the mosquitoes were offered the antiviral sugar bait solution for 4 days, followed by access to 10% sucrose solution until tissue collection.

Salivation and collection of mosquito tissue samples were performed at 7 and 14 days post infection (dpi) for the JNJ-A07-treated DENV group. Mosquito tissue samples were collected at 3, 7 and 14 dpi for the NHC-, MPV- and 4’FlU-treated CHIKV groups (Fig. 2).

#### Salivation and tissue dissection

At the selected time points, females were cold-anesthetized, wings and legs were removed from each mosquito using forceps. Forced salivation was then performed by inserting the proboscis of each mosquito in a tip filled with 20 µL of FBS for 1 hour. Next, the tip was collected, and the content was diluted in 40 µL of mosquito diluent (PBS supplemented with 20% FBS, 50 µg/mL penicillin-streptomycin, 50 µg/mL gentamicin, and 2.5 µg/mL Fungizone). Following the forced salivation, each mosquito head was separated from the body. The head, wings and legs of each mosquito were placed in homogenization tubes containing 2.8 mm ceramic beads (Precellys, Bertin Technologies, France) and 300 µL of mosquito diluent. The mosquito body was placed in a separate homogenization tube containing 300 µL of mosquito diluent.

All samples collected (saliva; body; head, wings, and legs) were stored at −80°C until further processing. The dissection forceps were disinfected in between samples by soaking in Virkon S (Lanxess, AG, Germany) followed by 70% ethanol to avoid cross-contamination.

#### Sample analysis

Mosquito bodies, heads, wings, and legs were homogenized using bead disruption at 7800 rpm for 1 min (Precellys Evolution, Bertin Technologies, France). The homogenate was spun down at 8000 rpm for 1 min. The supernatant was transferred to a 0.8 μm Vivaclear Mini filter (Sartorius, Germany) and then centrifuged at 10 000 rpm for 2 min. The filtered homogenate was used in end-point titration or focus-forming assays (FFA) to detect infectious virus.

### 10. End-point titration assay

Infectious virus titers in CHIKV-infected samples were determined by end-point titration on Vero cells. Cells were pre-seeded in a 96-well plate at a density of 25 000 cells/well in 180 µL of assay medium (2% FBS, 1% L-glutamine, 1% Sodium bicarbonate, 1% non-essential amino acids, 1% Penicillin-Streptomycin). The following day, infection was initiated by adding 20 µL of infectious sample to the first well, followed by a 10-fold dilution series. Plates were incubated for 3 days at 37 °C, after which cytopathogenic effect (CPE) was scored. Viral titers were calculated as TCID_50_/mL using the Reed and Muench method.

### 11. Focus forming assay (FFA)

Infectious DENV-2 titers in mosquito samples were quantified using FFA, as previously described [15]. Briefly, BHK cells were pre-seeded in a black 96-well tissue culture plate (1 × 10^5^ cells/well). The following day, cells were inoculated with mosquito samples for 2h at 37 °C. Following removal of the inoculum, cells were overlaid with 0.8% CMC diluted in supplemented RPMI medium and incubated for 3 days. Monolayers were subsequently fixed with 4% paraformaldehyde and permeabilized with 0.5% Triton X-100 in PBS. Cells were then stained with the anti-DENV complex antibody D3-2H2-9-21 [1:500 diluted in block buffer (3% bovine serum albumin, 0.2% Tween 20, and 2% FBS); MAB8705; Sigma-Aldrich], followed by an Alexa Fluor 594-conjugated secondary antibody (1:500 diluted in block buffer; A-11005; Invitrogen, Thermo Fisher Scientific) and DAPI (final concentration of 100 nM) counterstaining. Images were acquired using an Operetta CLS system (Revvity) and analyzed with Harmony software. DENV-2-infected cells were identified based on Alexa Fluor 594 positivity and foci used for quantification of infectious virus.

### 12. Data analysis

All figures and statistical analysis were made using GraphPad Prism v10.6.1 (GraphPad Software, San Diego, California USA). Infection rate (IR) was calculated as the proportion of blood-fed mosquitoes with infectious virus present in the body. Dissemination rate (DR) was the proportion of mosquitoes with a positive infection in the body that also had infectious virus in the head, wings, and legs. Transmission rate (TR) was the proportion of mosquitoes with a disseminated infection that also had infectious virus present in the saliva. IRs, DRs, and TRs were statistically compared using the Fisher’s exact test.

## Results

### The antiviral 4’fluorouridine (4’FlU) reduced CHIKV infection in *ex vivo* mosquito guts

NHC and JNJ-A07 were previously shown to inhibit the infection of CHIKV and DENV, respectively, in *ex vivo* mosquito guts [15, 16]. For 4’FIU however, no antiviral data in mosquito tissue were available yet. Therefore, the anti-CHIKV activity of 4’FlU was first examined in *ex vivo* cultured guts from *Ae. aegypti* to guide the selection of doses for subsequent *in vivo* administration. CHIKV-infected *ex vivo* guts were exposed to different concentrations of 4’FlU in the culture medium. In this model, treatment with 4’FlU resulted in a dose-dependent inhibition, with all tested concentrations showing a significant antiviral effect compared to the untreated control (DMSO). The lowest concentration (1 µM) resulted in a median reduction of 7.2-fold in infectious virus titer. Antiviral activity increased with rising 4’FlU concentrations, reaching a maximal reduction of approximately 178-fold at 200 µM (Fig. 3A).

**Fig. 3.**
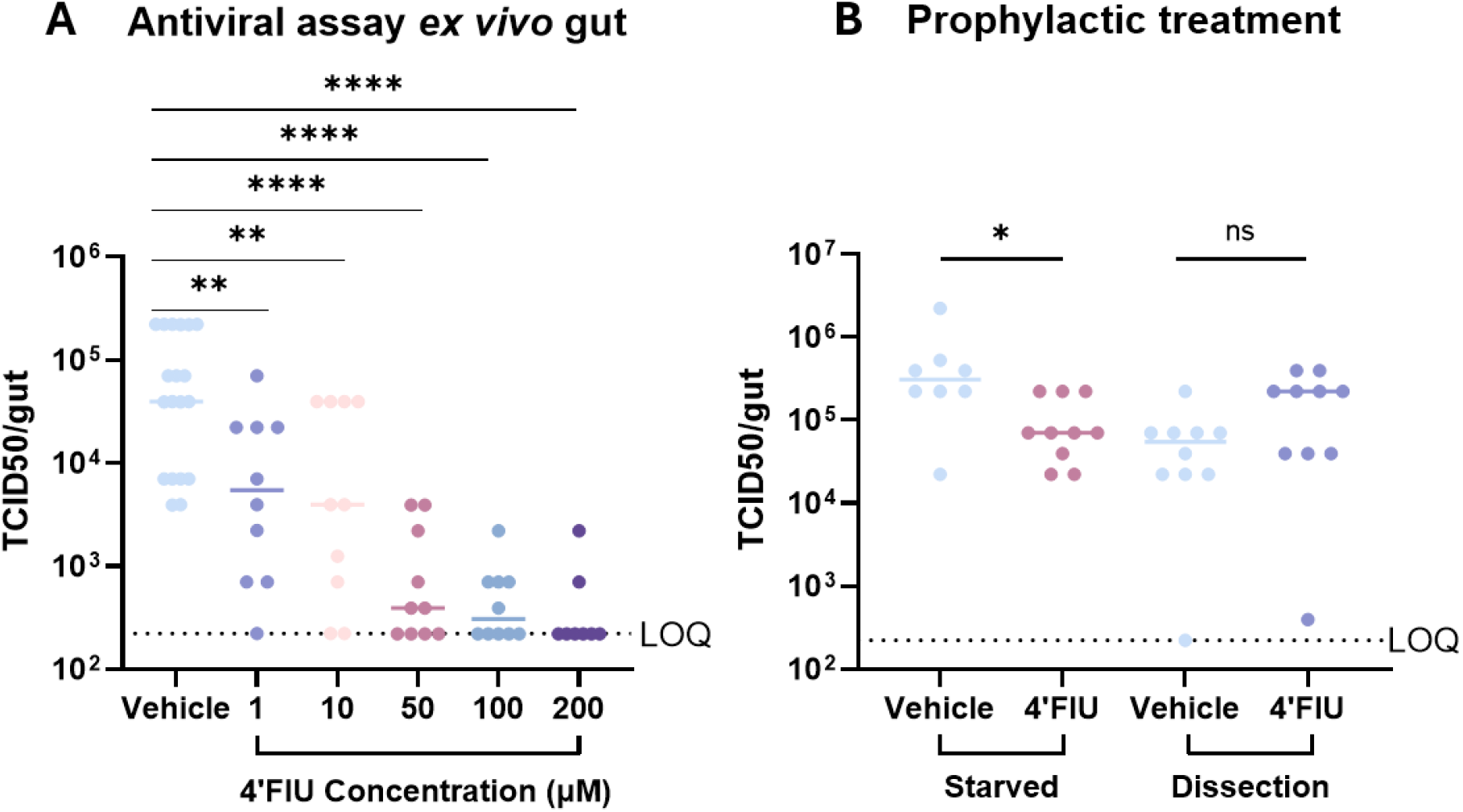
Antiviral activity of 4’FlU in *ex vivo* cultured mosquito guts. **(A)** Infectious viral titers in CHIKV-infected guts when incubated in the presence of 4’FlU for 3 days. **(B)** Infectious viral titers in CHIKV-infected guts following administration of 4’FlU (200 µM) via sugar bait for 24h, followed by a 24h starvation period before dissection (group 1) or by immediate dissection (group 2). Viral titers were measured at 3 dpi by end-point titration and are expressed as TCID50/gut. Statistical significance was evaluated with a non-parametric Mann-Whitney test (ns: not significant, *p<0.05, **p<0.01, ****p<0.0001). Each dot represents a mosquito gut. Horizontal lines represent the median value of each condition. Dotted lines depict the limit of quantification (LOQ) of the assay. Data shown corresponds to two independent experiments.

To investigate whether the antiviral effect of 4’FlU (200 µM) was maintained when administered through a sugar bait, mosquitoes were offered the sugar bait for a 24h-period and subsequently starved for 24h before the *ex vivo* experiment (group 1). This approach was used to determine whether the compound could exit the crop and reach the midgut, where its antiviral activity could be assessed. In the second group, mosquitoes were offered the sugar bait for 24h and were dissected immediately thereafter (group 2). In group 1, treatment with 4’FlU resulted in a significant but modest reduction in CHIKV infection in the *ex vivo* guts compared to the vehicle control (DMSO), with a median reduction of 4.4-fold(Fig. 3B). In contrast, viral loads in group 2 were similar between the 4’FlU treatment and control conditions.

### The olfactory attractiveness to AVSBs was not altered

Antiviral compounds may possess a repulsive odor that would negatively impact the effectiveness of the AVSBs. Therefore, a choice experiment was conducted in a partially closed system to test the mosquito attraction to the AVSB formulations. Mosquito landings on each AVSB-solution were recorded for a duration of 30 minutes. The presence of DMSO (vehicle control) alone did not affect mosquito attraction, as no significant difference in mosquito landings was observed between the vehicle-AVSB and a 10% sucrose solution (lacking DMSO) (Fig. 4A). Furthermore, no significant differences in mosquito landings were observed following the addition of JNJ-A07, MPV, NHC or 4’FlU to AVSBs as compared to the DMSO control baits (Fig. 4B-E). Additionally, none of the AVSB formulations exhibited reduced attractiveness relative to the control, with a preference index (PI) value ranging from 0.025 to 0.33.

**Fig. 4.**
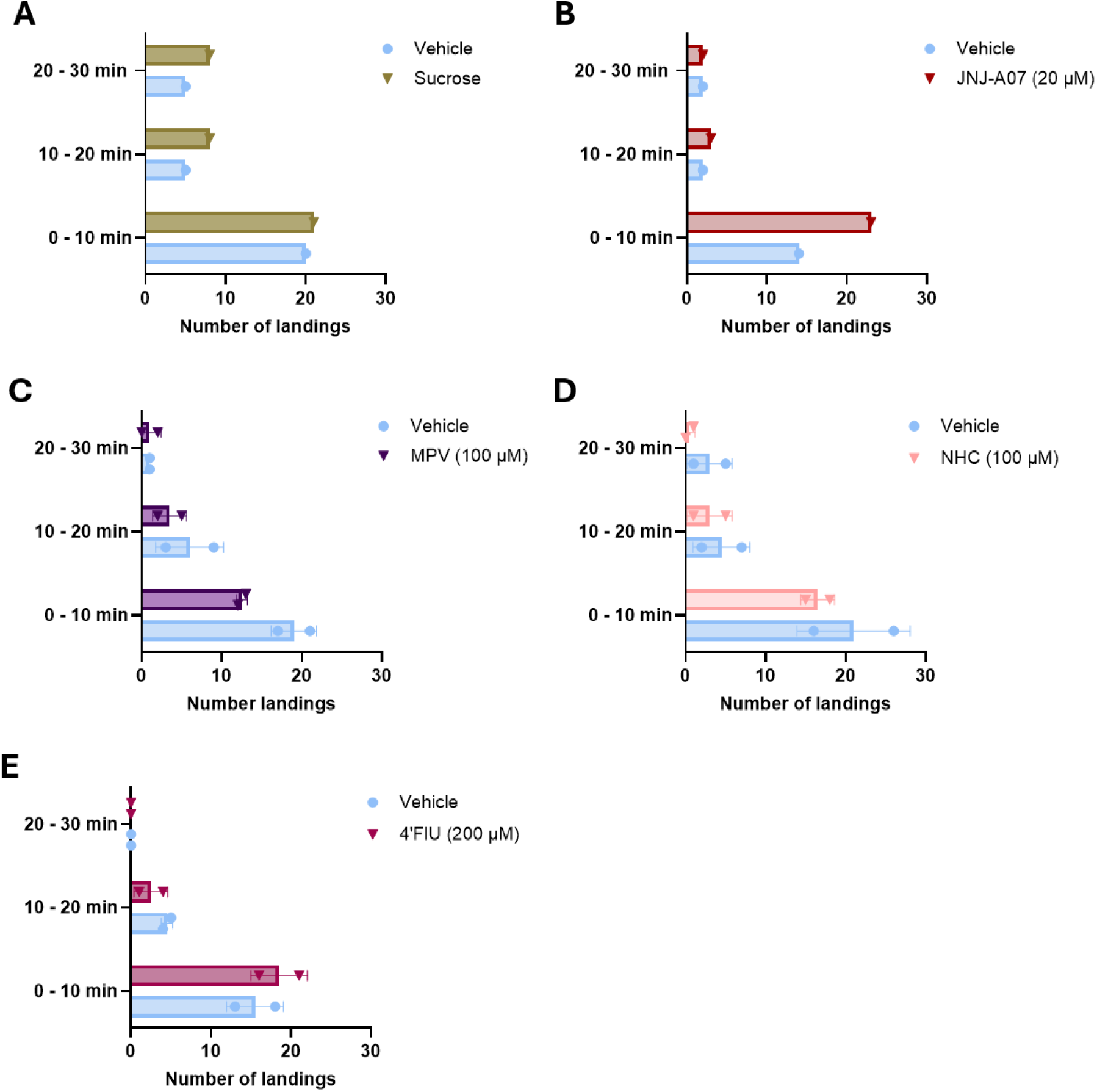
Mosquito olfactory attractiveness to antiviral sugar baits (AVSBs). Number of mosquito landings on **(A)** 10% sucrose (PI = 0.10), **(B)** JNJ-A07 AVSB (20µM; PI = 0.22), **(C)** MPV AVSB (100µM; PI = 0.18), **(D)** NHC AVSB (100µM; PI = 0.15), **(E)** 4’FlU AVSB (200µM; PI = 0.025), each compared with a corresponding 10% sucrose solution containing DMSO (vehicle control). Statistical significance was evaluated using the Fisher’s exact test. No significant differences were observed between treatments and controls (p>0.05). Data shown are from 2 independent experiments, except A and B. PI: Preference Index.

### AVSBs exposure had limited effects on mosquito longevity

To assess the effects of the anti-CHIKV compounds on the longevity of the mosquitoes, survival was monitored for 30 days following a 7-day exposure period to the AVSBs. Both male and female mosquitoes were observed (Fig. 5). NHC reduced female mosquito survival, resulting in a 30-day mortality rate of 80% compared with 47.5% in the corresponding control group. AVSBs containing MPV or 4’FlU had no significant effect on female mosquito survival relative to the control treatment. For male mosquitoes, none of the compounds significantly affected survival compared with their respective control groups, except for 4’FlU. Male mosquitoes exposed to 4’FlU exhibited a 30-day mortality rate of 70% compared with 30% in the control group, indicating reduced survival.**Female Male**

**Fig. 5.**
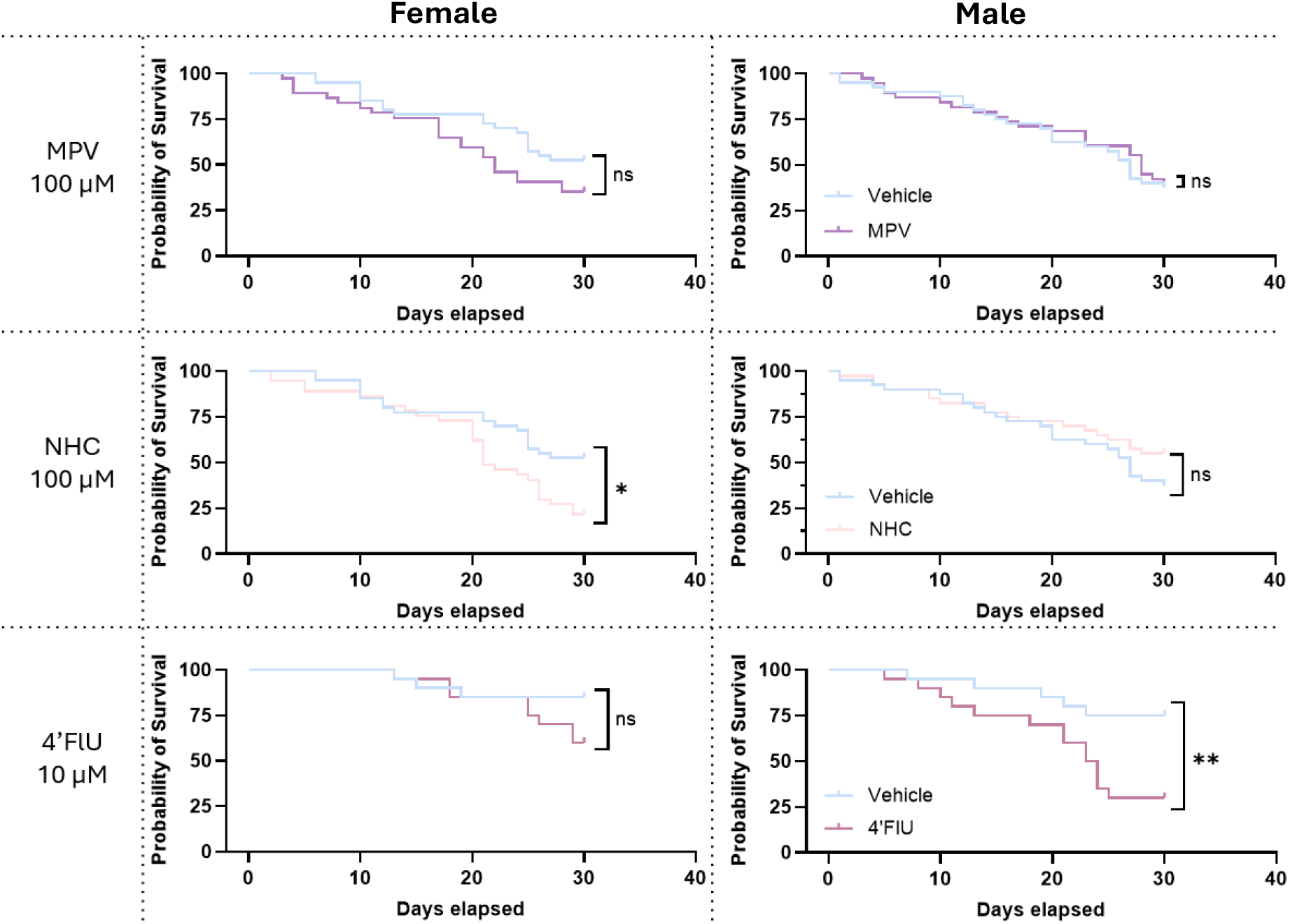
Survival of *Ae. aegypti* fed with an AVSB. Kaplan-Meier curves showing the proportion of surviving female mosquitoes (left panel) and male mosquitoes (right panel) monitored over 30 days after feeding on AVSB compared with vehicle AVSB (DMSO, control group). Statistical significance was assessed by two-sided log-rank (Mantel-Cox) test (*p<0.05, **p<0.01).

### Effect of AVSBs on fecundity and fertility

Female mosquitoes were exposed to AVSBs containing MPV (100 µM), NHC (100 µM), or 4’FlU (200 µM), or a DMSO (vehicle) sugar bait as a control, prior to being offered a blood meal. Four days after the blood meal, the females that received the MPV sugar bait had significantly fewer eggs in their ovaries than females in the control group. The mean (± SD) number of ovarian eggs was 75±23 and 95±22 in the MPV sugar meal and DMSO sugar bait, respectively (Fig. 6A). In contrast, sugar baits containing NHC and 4’FlU did not significantly affect the number of eggs developed in the ovaries. Despite the reduced number of ovarian eggs, females that received the MPV AVSB laid significantly more eggs than control mosquitoes, with a mean (± SD) of 95±23 and 67±39, respectively (Fig. 6B). Neither NHC nor 4’FlU significant affected egg-laying behavior of the female mosquitoes.

**Fig. 6.**
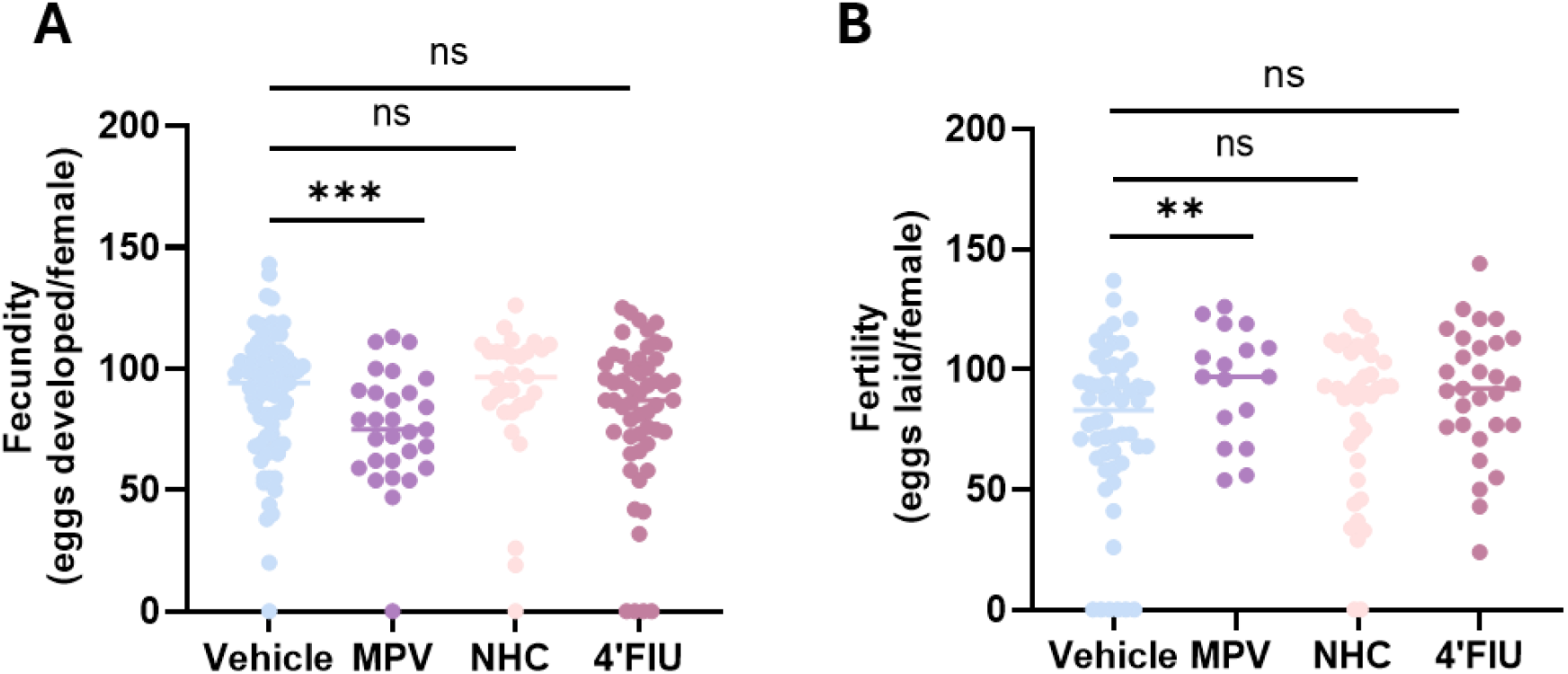
Effect of MPV-, NHC- and 4’FlU-AVSB on fecundity (A) and fertility (B) of *Ae. aegypti* mosquitoes. **(A)** Number of eggs developed inside the ovaries of mosquitoes that fed on a blood meal after exposure to MPV AVSB (100 µM; n=29), NHC AVSB (100 µM; n=30), 4’FlU AVSB (200 µM; n=54) and the vehicle AVSB (DMSO, n=73). **(B)** Number of eggs laid per female that fed on a blood meal after exposure to MPV AVSB (100 µM; n=17), NHC AVSB (100 µM; n=39), 4’FlU AVSB (200 µM; n=29) and the vehicle AVSB (DMSO; n=55). The blood meal for all conditions contained a final concentration of 0.5% DMSO. Statistical significance was evaluated with a non-parametric Mann-Whitney test (ns: not significant, **p<0.01, ***p<0.001). Each dot represents the data of an individual female mosquito. The horizontal lines indicate the median value per condition.

### The JNJ-A07 AVSB efficiently blocked DENV-2 transmission

In a previous study, JNJ-A07 had potent antiviral activity against DENV-2 infected *Ae. aegypti* mosquitoes when administered through a bloodmeal [15]. To investigate whether JNJ-A07 also exerted antiviral effects when delivered through a sugar meal, AVSBs containing JNJ-A07 (2 µM or 20 µM) were offered to female mosquitoes starting 3 days before DENV-2 infection and continuing until 4 days post infection (dpi) (Fig. 7A). At the 14 dpi endpoint, infection rates (IRs) in the vehicle control and 2 µM JNJ-A07 groups were 87% and 93%, respectively, whereas only 3% of mosquitoes in the 20 µM JNJ-A07 group were positive for DENV-2 infection (Fig. 7B). Disseminated rates (DRs), indicative of viral spread to secondary tissues, reached 100% in the vehicle control group. In comparison, DRs in the JNJ-A07-treated groups were 74% for the 2 µM concentration and 0% for the 20 µM concentration. Transmission rates (TRs) were 2%, 10%, and 0% in the vehicle control, 2 µM JNJ-A07, and 20 µM JNJ-A07 groups, respectively.

**Fig. 7.**
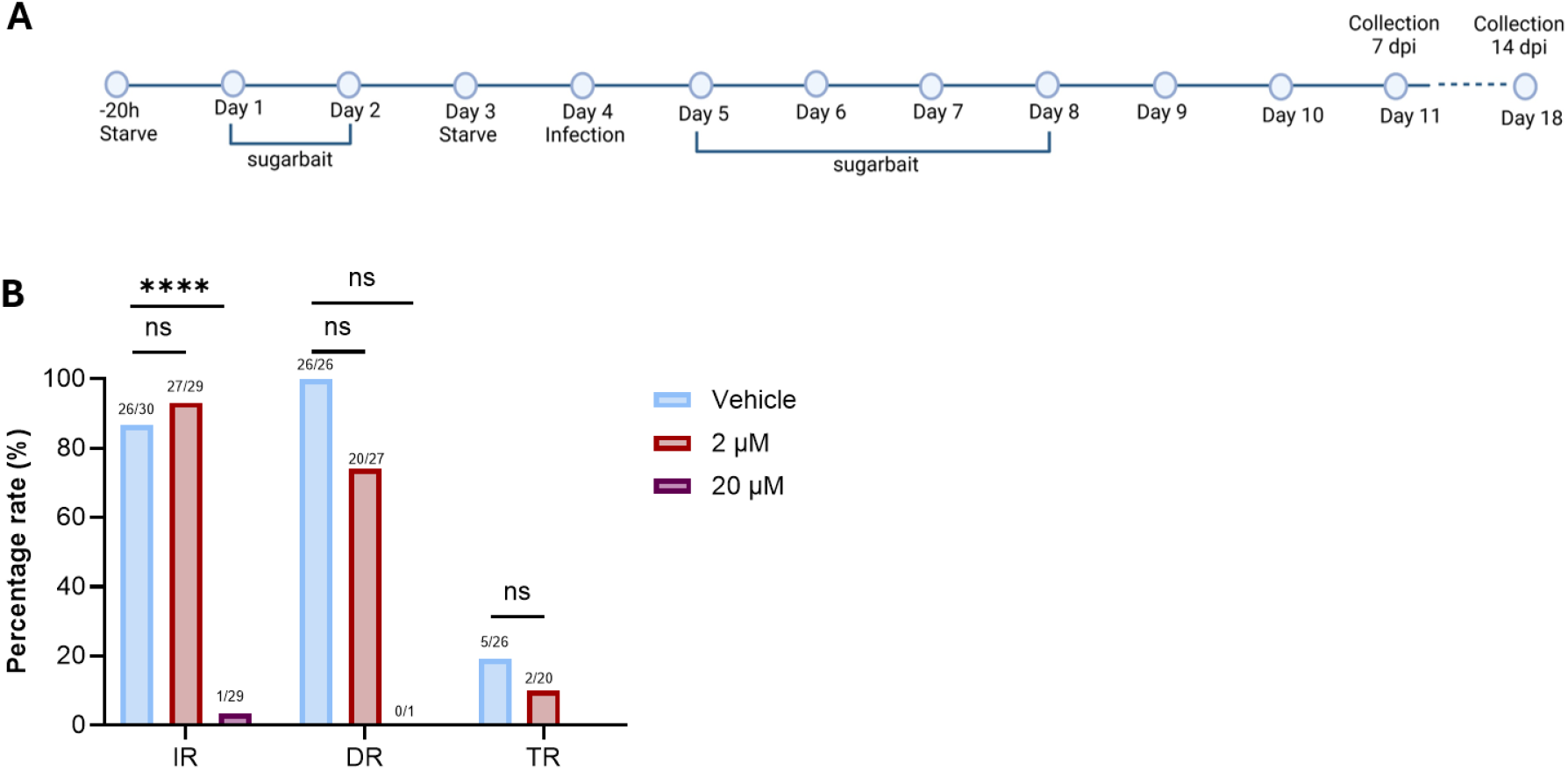
The antiviral effect of JNJ-A07 on the vector competence of *Ae. aegypti* mosquitoes infected with DENV-2. **(A)** Timeline depicting the design of the experiment, in which mosquitoes were offered the JNJ-A07 AVSB (2 µM or 20 µM) or vehicle AVSB (DMSO control group) for 3 days before and 4 days following a blood meal containing DENV-2. Mosquito tissues were collected at 14 dpi. Infectious virus was detected by foci forming assay. **(B)** DENV-2 infection (IR), dissemination (DR), and transmission rates (TR) for mosquitoes treated with JNJ-A07 AVSB (2 µM or 20 µM) compared to vehicle control (DMSO).

### The 4’FlU-AVSB efficiently blocked CHIKV infection in *Ae. aegypti*

Previous studies showed that NHC and MPV did not reduce CHIKV infection when administered to mosquitoes via a blood meal, despite antiviral activity in mosquito cells and *ex vivo* mosquito guts [16]. Here, we evaluated whether delivery through AVSBs could improve the antiviral efficacy of NHC and MPV *in vivo*. In addition, because 4′FlU potently inhibited CHIKV replication in the *ex vivo* mosquito gut model, its antiviral activity was also assessed *in vivo* following administration through AVSBs. To investigate whether these AVSBs exerted antiviral effects when delivered through a sugar meal, AVSBs containing NHC (100 µM), MPV (100 µM) and 4’FlU (200 µM) were offered to female mosquitoes starting 3 days before CHIKV infection and continuing until 4 days post infection (dpi) (Fig. 8A).

**Fig. 8.**
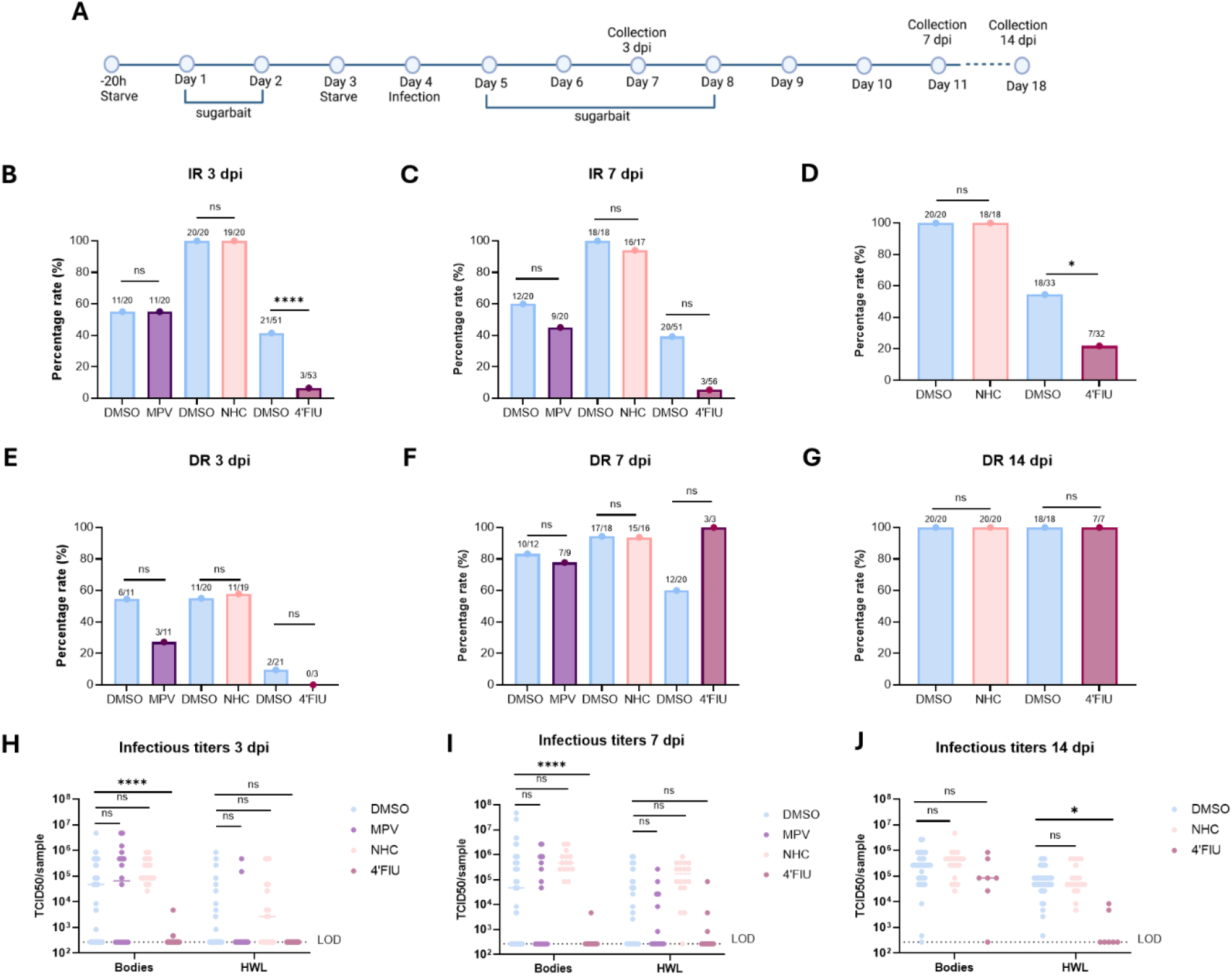
The antiviral effect of MPV, NHC, and 4’FlU on the vector competence of *Ae. aegypti* mosquitoes infected with CHIKV. **(A)** Timeline depicting the design of the experiment, in which mosquitoes were offered the MPV (100 µM), NHC (100 µM) or 4’FlU (200 µM) AVSB or vehicle AVSB (DMSO, control group) for 3 days before and 4 days following a blood meal containing CHIKV. Mosquito tissues were collected at 3, 7 and 14 dpi. Infectious virus was detected by end-point titrations. Infection rates at **(B)** 3 dpi, **(C)** 7 dpi and **(D)** 14 dpi. Dissemination rates at **(E)** 3 dpi, (**F)** 7 dpi and **(G)** 14 dpi Statistical significance determined using the Fisher’s exact test. ns, not significant (*p<0.05, **p<0.01, ***p<0.001, ****p < 0.0001). Viral titers in bodies and HWL at **(H)** 3 dpi, **(I)** 7 dpi and **(J)** 14 dpi. Statistical significance was determined using the Mann-Whitney test. ns, not significant; (*p<0.05, **p<0.01, ***p<0.001, ****p < 0.0001). Each dot represents the sample of an individual mosquito. Dotted lines depict the limit of detection (LOD) of the assay. HWL = head, wings, legs.

No significant differences in IRs or DRs were observed between the vehicle control, MPV, and NHC treatment groups at any of the investigated time points (3, 7, and 14 dpi) (Fig. 8B-G). In contrast, mosquitoes exposed to 4′FlU-AVSBs exhibited significantly lower infection rates than vehicle-treated mosquitoes at 3 and 14 dpi. Infection rates were reduced by approximately 35% an 33%, respectively. However, no significant differences in dissemination rates were observed between the 4′FlU and control groups. Consistent with these observations, viral titres in mosquito bodies were significantly reduced following exposure to 4′FlU-AVSBs, with reductions of 138-, 513-, and 355-fold at 3, 7, and 14 dpi, respectively (Fig. 8H-J). In addition, mosquitoes with disseminated infections exhibited significantly lower viral titres in head, wing, and leg tissues at 14 dpi, corresponding to a 2.3-fold reduction relative to the vehicle control group. In contrast, MPV- and NHC-containing AVSBs did not significantly affect viral titres at any time point.

## Discussion

Arboviruses such as DENV and CHIKV are major threats to global health; still effective antiviral therapies are lacking [22]. Although substantial mosquito control programs have been implemented in endemic regions, these strategies alone have not been sufficient to significantly reduce disease burden [23]. Consequently, alternative approaches that complement conventional vector control are needed. In this study, we evaluated the use of antiviral sugar baits (AVSBs) as a novel strategy for delivering antiviral compounds directly to mosquito vectors, with the aim of disrupting viral replication and reducing virus transmission to humans. Previous studies have demonstrated successful delivery of small molecules to mosquitoes through compound-spiked blood meals [15], tarsal exposure (e.g., insecticide-treated bed nets) [24], and attractive toxic sugar baits (ATSBs) [17]. However, each of these approaches has limitations. Blood meal-mediated delivery requires sufficient antiviral exposure of the vertebrate host, whereas tarsal exposure and ATSBs predominately rely on insecticidal compounds. While effective, insecticides may adversely affect non-target species, contribute to environmental contamination, and drive the emergence of resistance [25]. To address these limitations, we explored the possibility of replacing conventional insecticidal agents in sugar baits with antiviral compounds that target the pathogen rather than the mosquito itself. These AVSBs exploit the natural sugar-feeding behavior of mosquitoes to reduce viral loads within the vector and thereby potentially limit virus transmission.

To study the antiviral potential of AVSBs, we selected four antiviral compounds with distinct mechanisms of action and previously reported anti-arboviral activity: JNJ-A07, NHC, MPV and 4’FlU. These compounds had demonstrated antiviral efficacy in mammalian models and, in some cases, mosquito-derived models, making them suitable candidates for assessing whether antiviral activity could be achieved following sugar-mediated delivery.

An important aspect of this study was the use of an *ex vivo* mosquito gut model to guide subsequent *in vivo* experiments. This model served as an intermediate screening method for identifying biologically relevant antiviral compounds and concentrations prior to labour-intensive mosquito infection studies. However, while *in vitro* and *ex vivo* models provide valuable preliminary information regarding antiviral activity, efficacy ultimately must be confirmed in the context of the whole mosquito, where uptake, metabolism and tissue distribution may substantially influence compound activity. Concentrations selected for *in vivo* evaluation were therefore informed not only by *ex vivo* mosquito gut experiments but also by previous mosquito studies (based on compound-spiked blood meals). JNJ-A07 was evaluated at 2 µM and 20 µM, two concentrations that were highly effective against DENV in *ex vivo* gut cultures and *in vivo* after blood meal delivery [15], while NHC and MPV were each tested at 100 µM, concentrations that had previously demonstrated antiviral activity against CHIKV in mosquito cells and *ex vivo* gut cultures [16]. Since 4’FlU had not previously been evaluated in mosquito systems, additional *ex vivo* gut experiments were performed to identify a suitable concentration for subsequent *in vivo* experiments. Although a significant reduction of CHIKV replication was observed at a concentration as low as 1 µM, the highest tested concentration (200 µM) was selected for subsequent experiments because it resulted in the strongest inhibition of CHIKV replication (Fig. 3A).

While the *ex vivo* gut model was useful for identifying effective antiviral concentrations, it does not address whether compounds delivered through sugar feeding can reach the mosquito midgut in sufficient amounts to exert antiviral activity. We therefore next examined the fate of antiviral molecules following ingestion through AVSBs. Mosquitoes mainly rely on plant-derived sugars as an energy source, and following ingestion, the sugar meal is directed to the crop where it is stored and gradually released into the midgut [26]. In contrast, infectious blood meals bypass the crop and are delivered directly to the midgut, where arboviral infection is established [27]. Consequently, the success of AVSB-mediated antiviral delivery depends not only on antiviral potency, but also on the ability of compounds to leave the crop and accumulate within the midgut at concentrations sufficient to inhibit viral replication. Using 4’FlU as a model compound, we demonstrated that antiviral activity became apparent only after an additional starvation period following 4’FlU-AVSB prophylactic exposure (Fig. 3B). This delayed effect suggests that time is required for the compound to be released from the crop and reach the midgut. Together, these findings provide important proof-of-principle that antiviral compounds delivered through sugar feeding can access the target tissue relevant for arboviral replication and retain biological activity following their release.

The effectiveness of AVSBs depends not only on the antiviral activity of the delivered compounds, but also on their ability to remain attractive to mosquitoes. If the addition of antiviral compounds were to reduce bait attractiveness or exert repellent effects, mosquitoes might preferentially feed on alternative sugar sources in the environment, thereby limiting the effectiveness of the intervention. Encouragingly, none of the antiviral compounds tested altered mosquito attraction to the sugar bait (Fig. 4), suggesting that their incorporation did not interfere with the natural sugar-feeding behaviour of *Ae. aegypti*. This finding is important since sugar sources are abundant in natural environments and AVSBs must compete with these alternative feeding opportunities to achieve sufficient uptake. The absence of detectable repellence indicates that mosquitoes are unlikely to avoid the AVSB and may readily feed on it when deployed in the field, rather than preferentially seeking alternative sugar sources in the surrounding environment, supporting the feasibility of this delivery platform for transmission-blocking interventions. However, it will be necessary to confirm this in competition assays in which mosquitoes are provided access to both natural sugars and AVSBs.

In addition to maintaining attractiveness, an effective AVSB should ideally exert only limited effects on mosquito survival and reproductive fitness. Unlike conventional ATSBs, which aim to reduce mosquito populations through toxicity, AVSBs are intended to suppress pathogen replication while exerting minimal effects on the vector itself. Such an approach may impose lower selection pressure on mosquito populations and consequently reduce the likelihood of resistance development. Furthermore, minimising effects on mosquito survival could decrease unintended ecological impacts associated with mosquito-killing interventions. Overall, the antiviral compounds evaluated in this study generally exerted only modest and compound-specific effects on mosquito longevity (Fig. 5) and reproductive fitness (Fig. 6), suggesting that antiviral activity can be achieved without substantially altering mosquito traits.

Although several statistically significant differences in longevity, fecundity and fertility were observed, these effects were generally modest and did not indicate widespread toxicity associated with AVSB exposure. For example, MPV altered reproductive output by reducing female fecundity while simultaneously increasing fertility. Interestingly, although MPV reduced the number of mature ovarian eggs, treated females laid more eggs than control mosquitoes, suggesting that ovarian egg development and oviposition may be affected differently by MPV exposure. The biological basis of this observation remains unclear and warrants further investigation. In contrast, NHC and 4′FlU had no detectable effects on either reproductive parameter. Likewise, survival effects were dependent on both the antiviral compound and mosquito sex, with NHC reducing female longevity, and 4′FlU reducing male longevity (Fig. 5). Importantly, a pronounced decline in survival was observed across all treatment groups approximatively 20 days after the start of the experiment. Given that the mosquitoes were 5 to 7 days old at the onset of the study, this corresponds to an age of approximately 25 to 27 days, which closely matches the reported lifespan of *Ae. aegypti* (∼27 days) maintained at 28°C [28]. This observation suggests that much of the mortality observed during the latter stages of the experiments could be attributable to natural age-related mortality rather than treatment-specific toxicity. Collectively, these findings indicate that AVSB-delivered antivirals can be administered with limited effects on mosquito survival and reproductive fitness, supporting their development as transmission-blocking rather than mosquito-killing interventions.

Having established that AVSBs can deliver antivirals to the mosquito midgut while maintaining bait attractiveness and exerting only limited effects on mosquito fitness, we next evaluated whether antiviral delivery through this platform could effectively suppress arboviral infection *in vivo*. JNJ-A07-containing AVSBs (20 µM) significantly reduced DENV-2 infection rates at both 7 and 14 dpi and additionally reduced viral dissemination at 14 dpi (Fig. 7). These findings are consistent with previous studies demonstrating the antiviral activity of JNJ-A07 following delivery to mosquitoes through a blood meal, and therefore indicate that its antiviral efficacy is maintained following sugar-mediated administration. Interestingly, antiviral activity was only observed at the higher concentration (20 µM), whereas the lower concentration (2 µM) exerted little effect on DENV infection. This suggests that a threshold level of antiviral exposure may be required to achieve sustained suppression of DENV replication following sugar feeding. Because compounds delivered through AVSBs are first stored in the crop and subsequently released into the midgut before reaching other mosquito tissues, the lower concentration may not have achieved sufficient tissue levels to maintain antiviral activity throughout the course of infection. In contrast, the higher concentration likely ensured that adequate levels of JNJ-A07 were retained within the mosquito to effectively inhibit viral replication and subsequent dissemination. Given that dissemination beyond the midgut is a prerequisite for transmission, the significant reduction in disseminated infections is particularly encouraging and highlights the potential of AVSB-mediated antiviral delivery as a transmission-blocking strategy against DENV.

Following the successful reduction of DENV infection using JNJ-A07-containing AVSBs, we investigated whether this strategy could also be applied to CHIKV using NHC, MPV and 4′FlU. Interestingly, the antiviral efficacy of AVSBs proved to be highly compound-dependent (Fig. 8). While neither MPV- and NHC-containing AVSBs reduced CHIKV infection or dissemination, 4’FlU significantly reduced both infection rates and viral dissemination. The observed reductions in both infection and dissemination suggest that 4′FlU reached relevant mosquito tissues at concentrations sufficient to suppress CHIKV replication throughout the infection process. The absence of an antiviral effect for NHC- and MPV-containing AVSBs may be explained by the rapid excretion of these compounds from the mosquito. We previously showed that when NHC was delivered via a blood meal, approximately 90% of the NHC was no longer detectable 6 h post-feeding, as determined by ultra-high-pressure liquid chromatography [15]. Although the digestion and metabolization of a blood meal differ from those of a sugar meal, similarly rapid excretion rates may occur after NHC-AVSB ingestion, limiting the exposure time required for antiviral activity. In addition, MPV is a prodrug of NHC and requires intracellular activation to exert its antiviral effect. Following cellular uptake, MPV is hydrolyzed to NHC, which must subsequently undergo phosphorylation to its active triphosphate form. It is possible that one or more of these activation steps occurred inefficiently in mosquito tissues *in vivo*, resulting in insufficient levels of the active metabolite. Further studies investigating these processes in mosquitoes are needed to confirm this hypothesis. Collectively, these findings highlight the importance of considering mosquito-specific pharmacokinetics, tissue distribution, and metabolic activation when selecting antiviral compounds for AVSB-based interventions.

Despite the promising results obtained in this study, several questions remain before AVSBs can be considered as a potential arbovirus control strategy. In particular, the pharmacokinetics of antiviral compounds within mosquitoes, including their uptake, distribution, metabolism, and excretion, remain poorly understood and may strongly influence efficacy. Future studies should therefore investigate the characteristics associated with successful AVSB-mediated delivery. In addition, AVSBs should be evaluated under semi-field and field conditions, where environmental factors, competition with natural sugar sources, and mosquito feeding behaviour may influence bait uptake and antiviral activity. Determining how frequently mosquitoes encounter and consume AVSBs in nature will be critical for assessing their feasibility, scalability, and long-term effectiveness as a vector control strategy.

This study demonstrates that attractive antiviral sugar baits can successfully deliver antiviral compounds to *Ae. aegypti* mosquitoes and suppress arboviral infection following sugar-mediated uptake. AVSBs containing JNJ-A07 and 4′FlU significantly reduced DENV and CHIKV infection, respectively, providing proof-of-concept for a novel transmission-blocking strategy that targets the pathogen within the vector. Together, these findings show the potential of AVSBs as a complementary approach for arbovirus control and warrant further evaluation under semi-field and field conditions.

## Notes

### Competing Interest Statement

The authors have declared no competing interest.

### Funding

- VLAIO
- FWO: 1173726N

